# High-Throughput Screening Identifies Proteasome Inhibitors as Potentiators of Type III Interferon JAK-STAT Signaling and Antiviral Immunity

**DOI:** 10.64898/2026.10.12.758093

**Authors:** Jean-Patrick Parisien, Crystal O’Leary, Shima Shabani, Sara Fernandez-Dunne, Laura T. Huynh, Aleksandra K. Kurowska, Curt M. Horvath, Juan L. Mendoza

## Abstract

Type III interferon (IFN3, IFNλ) drives antiviral defense at epithelial barriers with a more favorable safety profile than Type I interferon (IFN1), but weaker signaling limits its therapeutic potential. To identify small molecules that potentiate IFN-JAK-STAT signaling, we developed a high-throughput live-cell assay measuring STAT1/STAT2 phosphorylation in A549 cells and screened ∼8,640 compounds. This identified proteasome inhibitors (MG132, delanzomib, ixazomib, oprozomib, bortezomib) as selective potentiators of IFN3-induced STAT activation, with minimal effect on IFN1 signaling. Proteasome inhibition enhanced IFN3 antiviral activity to levels approaching IFN1. Mechanistically, proteasome inhibition increased cell-surface IL10Rβ, the limiting IFN3 receptor subunit, thereby increasing receptor JAK-STAT signaling. It did not alter IFN1 receptor levels but protected IFNAR2 from late ligand-induced downregulation. These findings establish receptor subunit availability as a rate-limiting, druggable node governing IFN3 signaling strength and demonstrate a scalable platform for identifying cytokine-selective signaling potentiators, with implications for mucosal antiviral therapeutics.

## INTRODUCTION

Antiviral immunity relies on the actions of interferon (IFN) family cytokines^1^. Type I IFN (IFN1, e.g., IFNα, IFNβ, IFNω) is well known to activate JAK-STAT signaling to induce target gene expression and antiviral responses in virtually all nucleated cells throughout an infected organism^2^. Epithelial barrier surfaces are buttressed by an additional line of host defense against respiratory, enteric, and mucosal pathogens through the actions of Type III IFNs (IFN3, e.g., IFNλ1/IL29, IFNλ2/IL28A, IFNλ3/IL28B, and IFNλ4)^3–5^. IFN3 is produced and signals at these barriers, using a heterodimeric receptor comprised of IFNλR1 (IL-28RA) and IL-10Rβ subunits, which is expressed predominantly on epithelial cells and a subset of immune cells^6^, in contrast to the ubiquitously expressed type I interferon receptor (IFNAR1/IFNAR2)^7,8^. The barrier-restricted IFN3 receptor expression confers a key therapeutic advantage: IFN3 elicits compartmentalized antiviral immunity without the broad systemic inflammation and associated toxicities that are known to limit the clinical utility of IFN1^9–11^.

The two IFN systems use distinct ligands and receptors; both IFN receptor signaling complexes converge intracellularly to activate JAK-STAT signaling to assemble the ISGF3 transcription regulator that contains a heterodimer of tyrosine-phosphorylated STAT1 (pSTAT1) and STAT2 (pSTAT2)^12–15^. However, IFN1 and IFN3 responses are quantitatively and kinetically distinct^16^. IFN3 induces weaker but more sustained pSTAT1 and pSTAT2 activity than IFN1, resulting in delayed, weaker, but prolonged expression of interferon-stimulated genes (ISGs) ^17,18^. This reduced signaling amplitude is thought to reflect intrinsic negative regulatory mechanisms or structural constraints on receptor-proximal JAK kinase activity and STAT activation that limit the maximal antiviral potency of IFN3 ^19^.

Small molecule modulation of cytokine-JAK-STAT signaling pathways is clinically validated: JAK inhibitors (jakinibs) have achieved widespread use in inflammatory diseases, demonstrating that receptor-proximal signaling nodes are tractable drug targets ^20,21^. One challenge is that jakinibs have varying levels of specificity for individual JAK family members and that inhibition of one JAK impacts dozens of cytokine pathways ^22^. Furthermore, compounds that selectively enhance rather than inhibit JAK-STAT signaling, particularly in a cytokine-specific manner, are desirable but far more challenging to identify. For example, a small molecule capable of boosting IFN3 signaling amplitude and antiviral activity would represent a novel tool for mucosal antiviral defenses, potentially bypassing the toxicity constraints that have limited systemic IFN1-based therapies ^9,10^.

We aimed to find small molecules that increase the signaling strength of IFN3s without the need for gene induction and protein translation that occur on an hours-to-days time scale typical of phenotypic screens and which may lead to identification of off-target compounds. Thus, we hypothesized that by monitoring pSTAT1/2 on the minutes-to-hour time scale, this approach would identify such compounds; we could develop a quantitative, cell-based high-throughput screening assay that measures pSTAT1 and pSTAT2 as a direct readout of IFN JAK-STAT signal transduction. Measuring pSTAT1/2 after 30 minutes of IFN stimulation would reduce false positives and favor compounds that act at a receptor-proximal step before signal amplification through transcriptional cascades. Simultaneous analysis of IFN1 and IFN3 responses on the same plate enables our platform to identify different classes of compounds that could potentiate the signaling of 1) IFN1-alone, 2) IFN3-alone, and 3) both IFN1 and IFN3.

Here we report the development and validation of this screening platform, and the results of a pilot screen against ∼8,640 compounds. One group of identified IFN signaling potentiators shared proteasome inhibitor activity, and we show they effectively enhance IFN3-mediated pSTAT1 and pSTAT2 activity and increase IFN3 potency in mediating antiviral activity. Mechanistic investigation reveals that proteasome inhibition dramatically increases the cell surface level of IL-10Rβ to increase sensitivity to IFN3 stimulation. In contrast, IFN1 receptors are only slightly increased prior to stimulation, but ligand-mediated degradation of IFNAR2 is blocked, enabling prolonged signaling to pSTAT1 and pSTAT2. These findings validate the screening approach and define a new regulatory node in IFN signal transduction with implications for IFN3 barrier antiviral immunity and therapeutic development.

## RESULTS

### Development and validation of a cell-based high-throughput assay for IFN signaling

To create a scalable platform for identifying small-molecule modulators of interferon signaling, we sought a quantitative, cell-autonomous readout of JAK-STAT pathway activation. Stimulation of A549 human lung epithelial cells with either IFN1 or IFN3 results in rapid, specific pSTAT1 and pSTAT2, detectable by immunoblot within 30 minutes and reaching peak levels within 1 hour of stimulation **(Fig. 1A, B)**. This early kinetic response reflects receptor-proximal JAK kinase activity and precedes transcriptional amplification, making it a stringent and direct measure of pathway activation.

**Figure 1.**
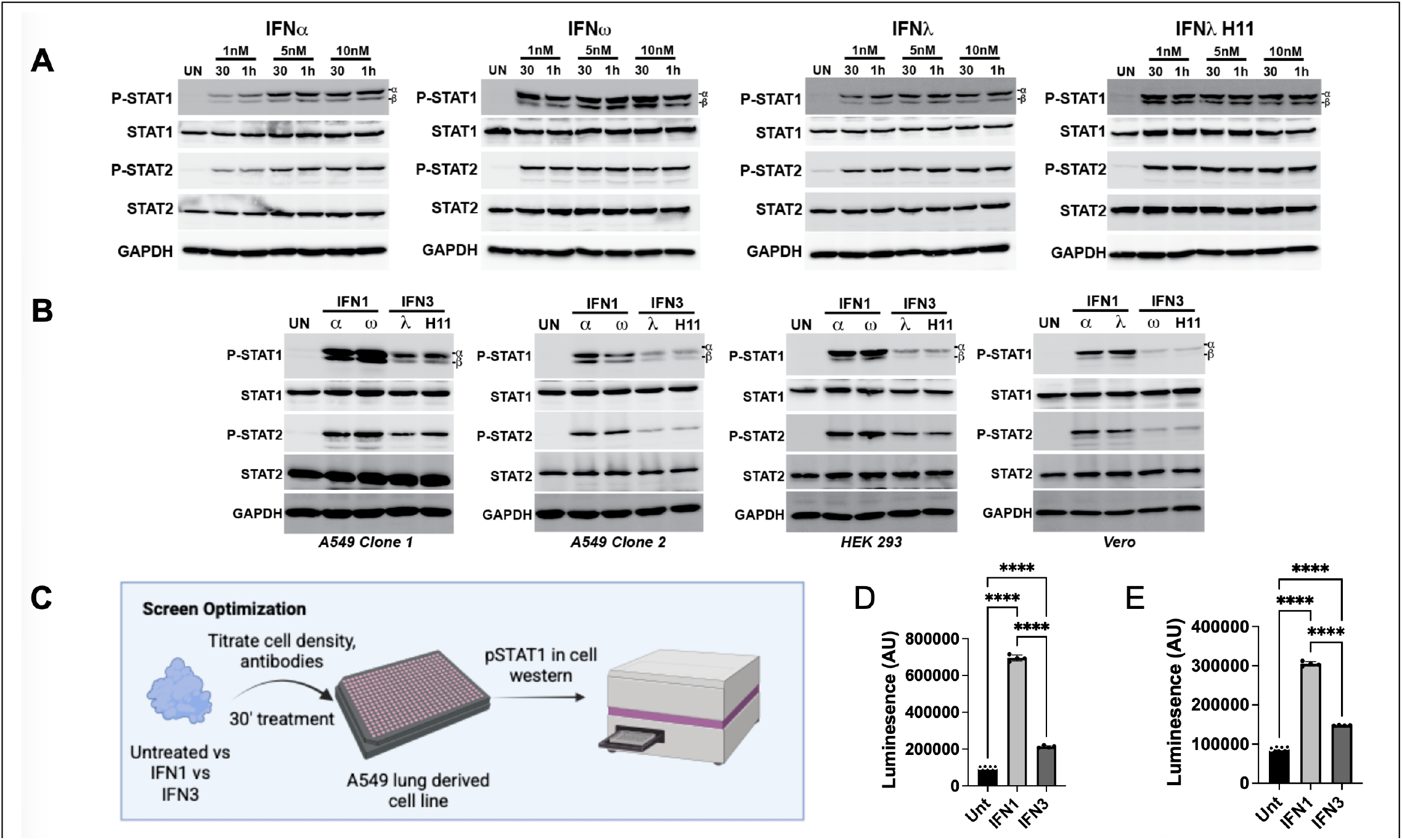
Design and validation of the cell-based high-throughput IFN-JAK-STAT signaling assay. **A)** A549 cells were stimulated with IFNα, IFNω, IFN3, or IFN3 H11, or left unstimulated at varying concentrations and times. **B)** The difference in signaling intensity by Western blot between IFN1s and IFN3s is observed in different cell lines: A549 Clones 1 and 2, HEK293, and Vero. **C)** Workflow schematic for compound library screening using the in-cell western (ICW) pSTAT1/pSTAT2 platform. In the optimized screen, **D)** pSTAT1 and **E)** pSTAT2 were quantified in 384-well formats under optimized screening conditions at a cell density of 10,000 cells/well. N=4 wells per condition, shown as the mean ± SD. All conditions differ significantly, indicating optimal conditions for a high-throughput screen.

To adapt this readout for a high-throughput format, we utilized and optimized an in-cell western (ICW) assay, a cell-based ELISA performed in fixed, permeabilized cells in multi-well plates that combines the target specificity of immunoblotting with the quantitative throughput of ELISA. This approach enables rapid, reproducible, quantitative measurement of pSTAT1 and pSTAT2 across hundreds of wells without cell lysis or gel electrophoresis. We systematically optimized cell density, IFN concentration, stimulation time, and detection parameters for multiwell format compatible with high-throughput screens **(Fig. 1C)**.

Consistent with immunoblot data, IFN1 induced stronger pSTAT1 and pSTAT2 signals than IFN3 in the ICW format, with both cytokines producing reliable signals above background across all tested conditions **(Fig. 1D, 1E)**. To assess assay quality, we calculated the Z’ factor, a statistical measure of assay robustness that accounts for signal dynamic range and variability, using wells treated with IFN alone (positive control) versus unstimulated wells (negative control). The assay achieved a Z’ factor ≥ 0.65 across replicate pilot experiments, indicating excellent assay performance suitable for primary high-throughput screening (Z’ > 0.5 is considered acceptable; > 0.5–1.0 excellent). The simultaneous measurement of IFN1 and IFN3 responses in the same experiment provides an internal counter-screen that controls for non-specific or cytotoxic effects and enables identification of different classes of potentiating compounds.

### Screening ∼8,640 compounds identifies modulators of IFN signaling

To demonstrate the capacity of this assay to identify small-molecule modulators of IFN signaling, we conducted a screen using six compound libraries available through the HTAL, representing approximately 8,640 structurally diverse and bioactive small molecules **(Fig. 2A)**. Compounds were added to A549 cells at a single screening concentration for 1 h before a 30 min stimulation with either IFN1 or IFN3 H11, an engineered high-affinity IFN3^19^, IFN3 H11 has the same signaling maximum as the wild-type IFN3 but has a higher sensitivity therefore requiring much less protein for the screen. On-plate controls included unstimulated/no-compound wells and IFN-only wells (no compounds). Following IFN stimulation, cells were processed for pSTAT1 by ICW. Luminescent values for each well were normalized to on-plate controls, and compound effects were expressed as percent modulation relative to the IFN1- or IFN3-only responses.

**Figure 2.**
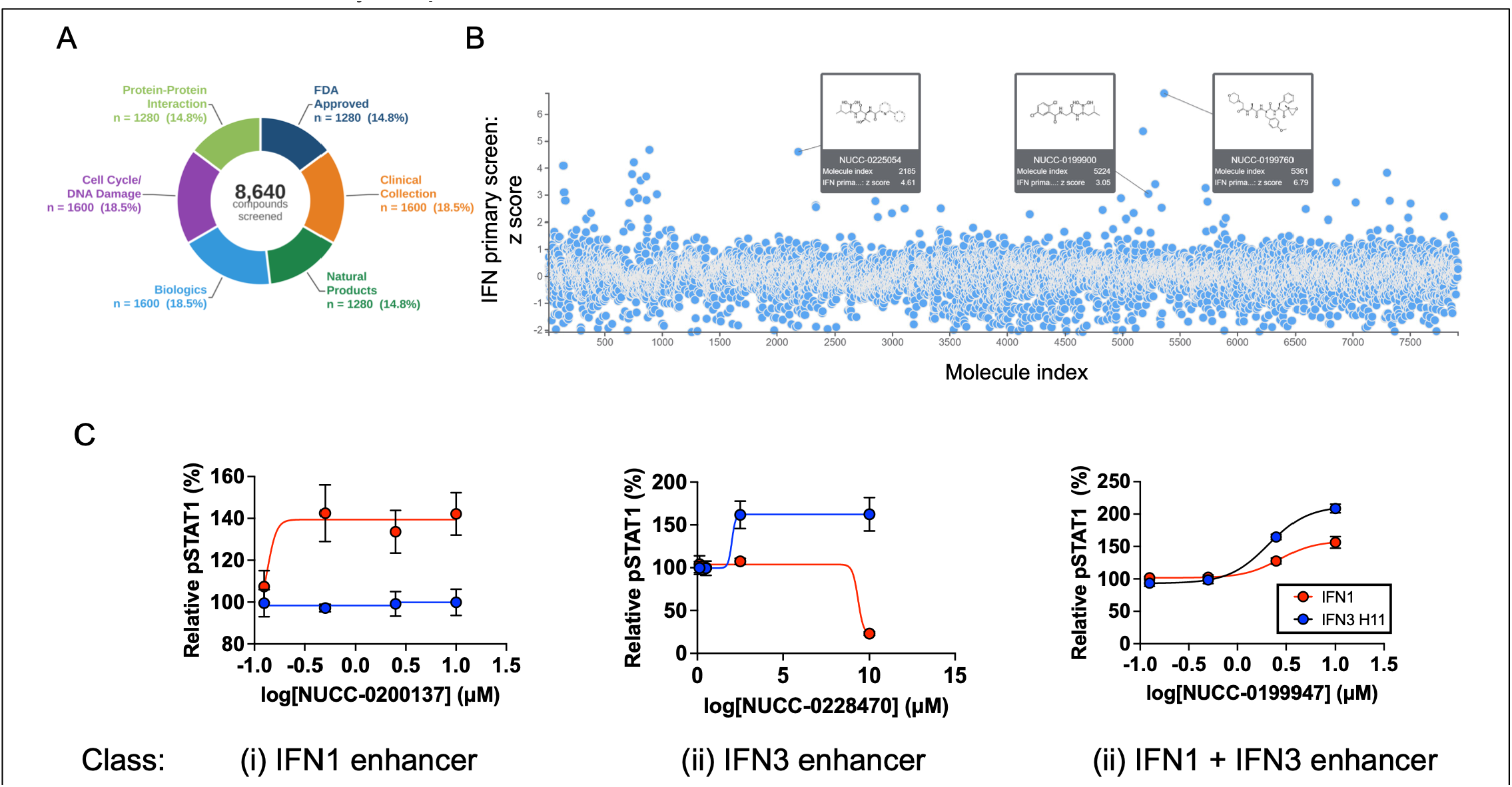
Screen for modulators of IFN signaling identifies three classes of activators. **A.)** Pie chart indicates the compound libraries used for the study. **B)** Dot plot of pSTAT1 modulation (expressed as % of IFN-only control) for all screened compounds, stratified by IFN type. Each dot represents one compound. Dashed lines indicate hit thresholds. Known JAK inhibitors are indicated. (C) Representative dose-response curves for compounds illustrating three categories: IFN3-selective enhancer (left), IFN1-selective enhancer (center), and dual enhancer (right). IFN1 responses are shown in red; IFN3 responses in blue.

The distribution of compound effects across the library is shown in **Fig. 2B**. As expected, most compounds had no significant effect on pSTAT1. A subset of compounds reduced pSTAT1 levels and included known JAK inhibitors and cytotoxic compounds. Although our primary interest was compounds potentiating IFN3 signaling, compounds that increased pSTAT1 over their matched IFN control (IFN1 or IFN3) were flagged as candidate enhancers and subjected to further validation and analysis.

Screen hits were ranked by magnitude of pSTAT1 enhancement and classified by their selectivity for IFN1 vs. IFN3: (i) enhanced IFN1, (ii) enhanced IFN3, or (iii) enhanced IFN1 and IFN3. Validation by performing dose-response assays for pSTAT1 signaling was performed on all three classes of compounds identified **(Fig. 2C)**. Among these hits, we identified a group of compounds that shared proteasome inhibitor activity, namely, MG132, Delanzomib, Ixazomib, Oprozomib; Bortezomib, which were selected for further analysis in dose-response experiments. Their shared target specificity and consistent strengthening of IFN3 signaling to pSTAT1 prompted detailed downstream characterization.

### Proteasome inhibitors enhance IFN3-mediated STAT1 and STAT2 phosphorylation

To validate the screen results and characterize the kinetics of enhancement by proteasome inhibitors, we performed time-course stimulation experiments. A549 cells were pre-treated with vehicle or MG132 1 h prior to stimulation with IFN1, IFN3, or IFN3 H11, and cell lysates were analyzed by immunoblot for pSTAT1 (Y701), pSTAT2 (Y689), and total STAT1/2 **(Fig. 3)**.

**Figure 3.**
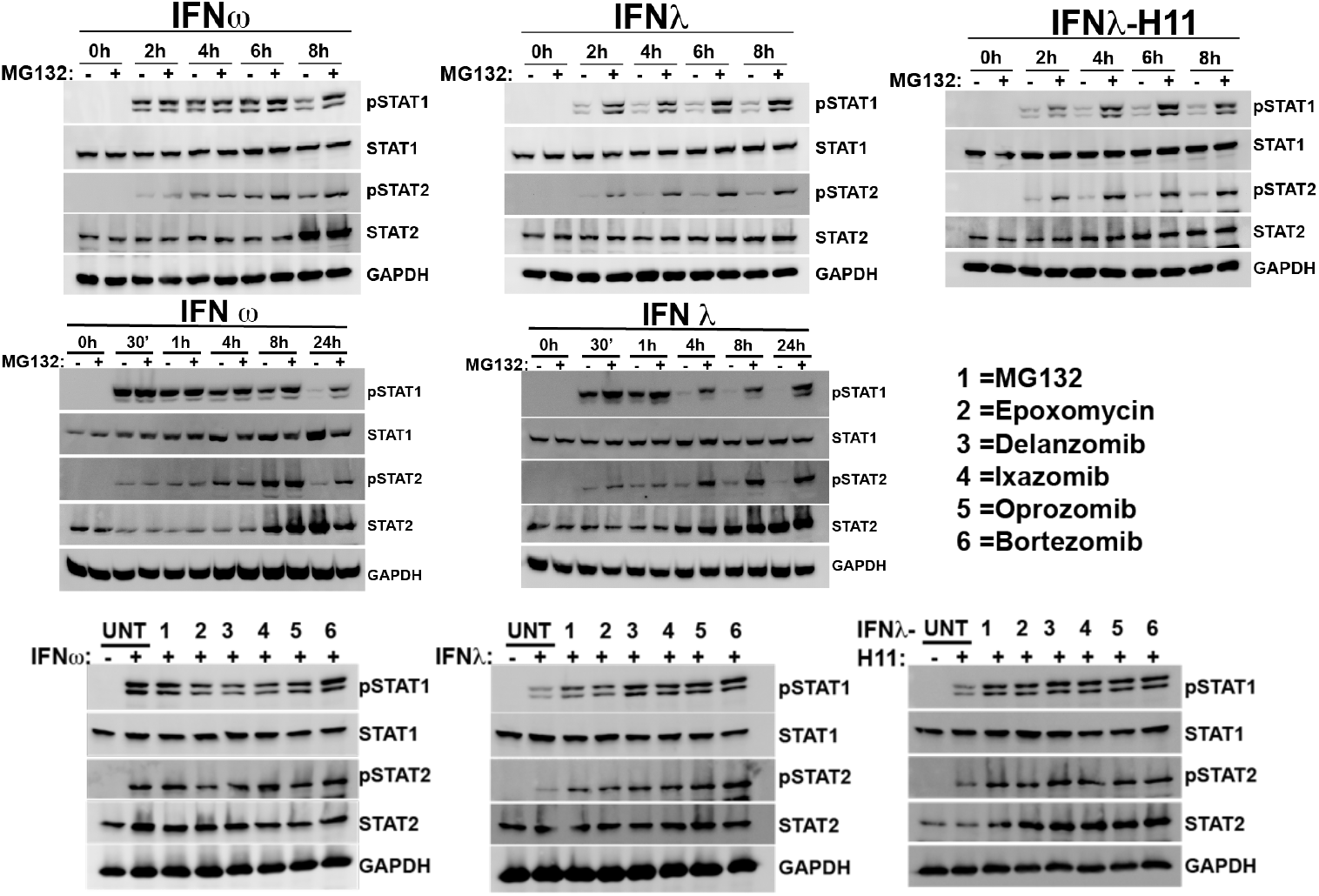
Proteasome inhibitors enhance IFN3 signaling to STAT1 and STAT2. (A) A549 cells were stimulated with IFN1, IFN3, or IFN3 H11 for the indicated times in the presence of vehicle (™) or MG132 (+). Lysates were analyzed by immunoblot for pSTAT1 (Y701), pSTAT2 (Y689), and total STAT1 (loading control). (B) Similar time-course experiment performed in HEK293T cells. (C) List of compounds used in panel D. (D) Single-time-point (4h) stimulation of A549 cells with IFN1, IFN3, or IFN3 H11 in the presence of vehicle or indicated proteasome inhibitors. Immunoblot for pSTAT1, pSTAT2, and STAT1. Data representative of n≥3 independent experiments.

MG132 had a minimal effect on IFN1-induced pSTAT1 or pSTAT2 as measured by western blot throughout most of the time course, although a clear effect was observed at the 8-hour time point in A549 cells. This suggests MG132 prevented the downregulation of STAT phosphorylation for IFN1-stimulated cells **(Fig. 3A)**.

In the first time-course experiment, in contrast to IFN1, MG132 markedly increased STAT1 and STAT2 activation by IFN3, observed at the earliest time point of 2 h post-stimulation in A549 cells. This signaling enhancement was sustained at all subsequent time points through 8 hours. To assess generality and rule out a cell line-dependent effect, we repeated the experiment in HEK293T cells, also an IFN1/3-responsive cell line **(Fig. 3B)**. Again, MG132 robustly enhanced IFN3-mediated pSTAT1 and pSTAT2 from 30 minutes through 24 hours. In addition, these experiments further demonstrated that the opposed ligand-induced attenuation observed for IFN1 signaling was also not cell-line dependent.

To confirm that these findings are shared across the other related compounds, we performed a single-time-point (4 h) comparison in A549 cells alongside a control structurally distinct proteasome inhibitor, epoxomycin. All the compounds significantly increased IFN3-mediated pSTAT1 and pSTAT2 while having little or no effect on IFN1 signaling at 4 h **(Fig. 3)**. These data establish proteasome inhibitors as a family of reproducible potentiators of IFN3-mediated STAT activation, with an onset of action within 30 minutes of cytokine stimulation.

### Proteasome inhibitors potentiate the antiviral activity of IFN3

The canonical function of IFN3-induced JAK-STAT signaling is to establish an antiviral state at barrier sites through the induction of IFN-stimulated genes that confer resistance to viral replication ^13^. To determine whether the enhanced STAT phosphorylation induced by proteasome inhibition translates to augmented antiviral biological activity, we used a reporter virus infection assay **(Fig. 4)**. A549 cells were treated with IFN1 or IFN3 with or without MG132 for 4 h to establish the antiviral state, then challenged with vesicular stomatitis virus encoding GFP (VSV-GFP) ^23^, a replication-competent prototypic IFN-sensitive negative-strand RNA virus **(Fig. 4)**.

**Figure 4.**
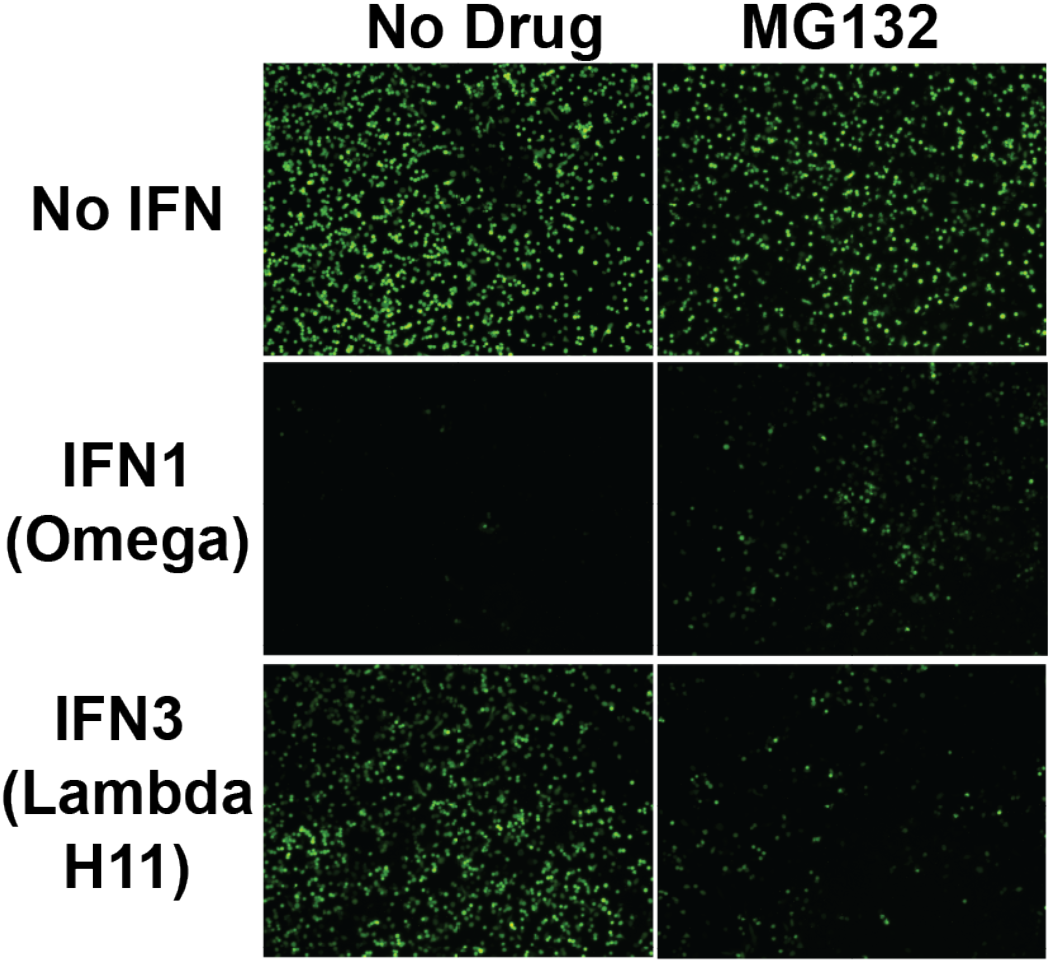
MG132 enhances the antiviral activity of IFN3. A549 cells treated with IFNs with or without MG132, as indicated, were challenged with VSV-GFP infection. IFN1 generates a strong antiviral response, but IFN3 only partially protects cells compared to No IFN. The addition of MG132 boosts IFN3 activity, reducing the VSV-GFP signal and slightly decreases IFN1 activity. Representative of N>3 replicates; similar results were observed for all proteasome inhibitors.

In the absence of MG132, IFN1 induced a strong antiviral state, nearly completely suppressing VSV replication as evidenced by near-absent GFP fluorescence. IFN3 reduced virus replication compared to unstimulated controls, but as expected, the antiviral activity was substantially weaker than IFN1, leaving a significant GFP-positive cell population. MG132 treatment did not affect VSV-GFP replication in the absence of IFN, confirming that the compound itself is not directly antiviral. Importantly, MG132 did not further enhance the already robust IFN1 antiviral response (and even showed a slight reduction). In contrast, MG132 dramatically potentiated IFN3 antiviral activity, reducing GFP-positive cells to levels approaching those achieved by IFN1 alone. This enhancement was reproducible across ≥3 independent experiments and was observed for all proteasome inhibitors tested, consistent with the enhanced signaling of IFN3-treated cells **(Fig. 4)**.

These results demonstrate that boosting IFN3-mediated STAT phosphorylation sufficiently translates into increased host defense against a viral challenge. As expected, IFN1 antiviral activity approaches maximal potency^19^, and despite prolonged signaling kinetics, it is not enhanced by MG132 in this biological assay.

### Mechanistic investigation reveals a specific regulatory target within the IFN/JAK/STAT pathway

MG132 potentiates IFN3 function in both signaling and antiviral assays, and enhancement of pSTAT1/2 is observed within 30 min of stimulation. The ability to boost IFN3 signaling on this fast time scale suggests the effects were likely acting at the receptor-proximal signaling level rather than processes like gene activation and protein translation that require hours-to-days to take effect. The ubiquitin–proteasome system governs the abundance and action of many cytokine receptors and controls ligand-induced degradation ^24–26^ . Thus, we performed flow cytometry-based experiments to measure the cell surface abundance of each subunit of the IFN1 (IFNAR1 and IFNAR2) and IFN3 receptors (IFNλR1 and IL10Rβ) **(Fig. 5A)**.

**Figure 5.**
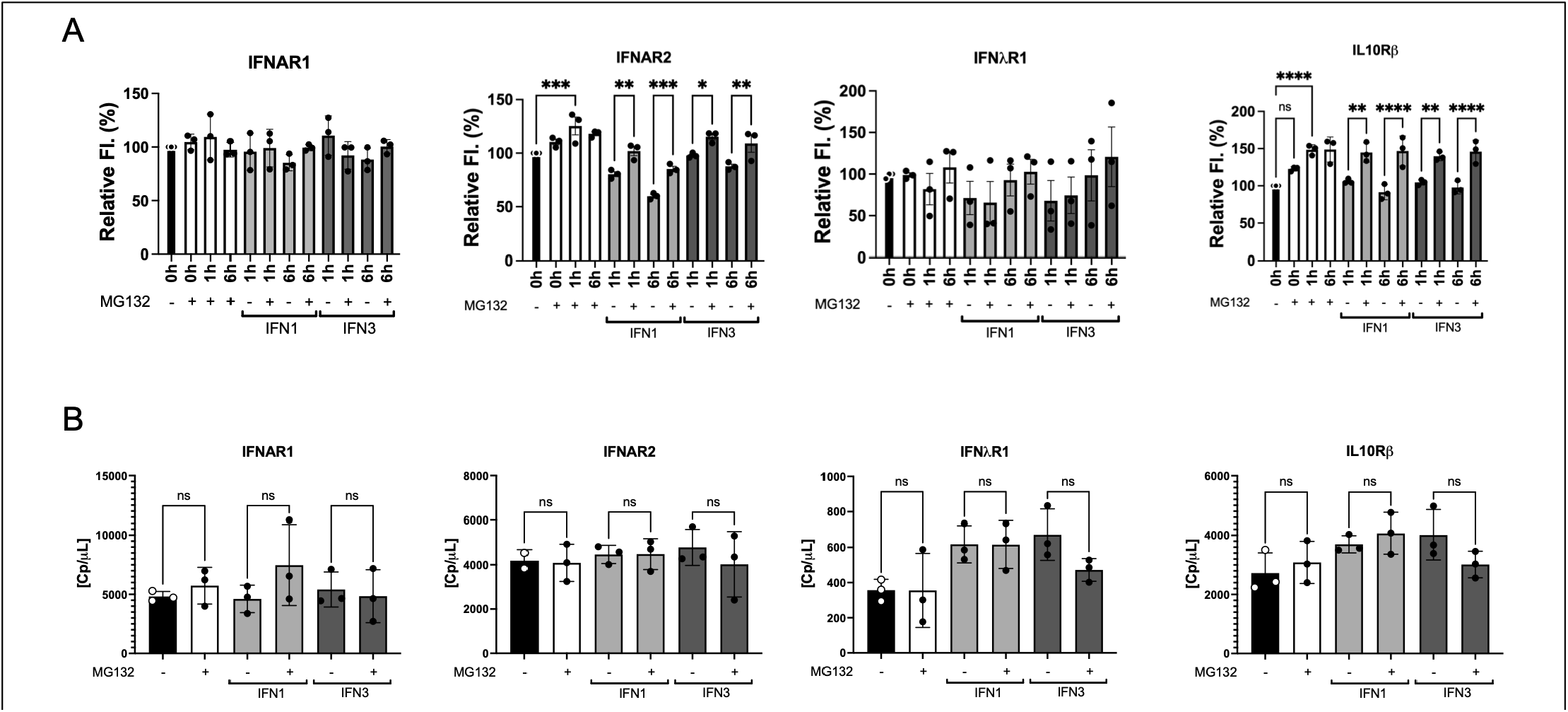
Proteasome Inhibition increases the IFNAR2 and IL10Rβ surface expression at the protein, not RNA, levels. **A)** A549 cells were pre-incubated with MG132 for 1 hour, then stimulated with IFN1 or IFN3 H11 for 0, 1, or 6 hours. Cells were stained with the respective antibody staining for IFNAR1, IFNAR2, IFNLR1, or IL10RB, and fluorescence was measured by flow cytometry. MG132 increases the expression of IFNAR2 and IL10RB within 1 hour and is sustained at 6 hours. **B)** A549 cells were pretreated with +/-CPD1 for 1 hour, after which no IFN or IFN1 or IFN3 was added. At 24 hours, the cells were harvested, RNA extracted, and converted to cDNA for dPCR quantification.

A549 cells were pre-incubated with MG132 for 1 hour, then stimulated with IFN1, IFN3, or left unstimulated, and surface expression of each subunit was quantified by flow cytometry with specific antibodies at 0, 1 h, and 6 h. MG132 produced a receptor-specific effect. Neither IFNAR1 nor IFNλR1 surface levels were significantly altered by MG132 under any condition or time point **(Fig. 5A)**.

In contrast, both IFNAR2 and IL10Rβ surface abundance were affected by MG132. A small increase in surface staining of IFNAR2 was observed in the MG132-only treated control, leading to a maximum of ∼1.25-fold by 1 h (P < 0.001) that is sustained at 6 h. Notably, IFN1 stimulation (without MG132) reduced surface IFNAR2 over time as expected by ligand-induced downregulation. MG132 restored IFNAR2 levels to or near those of unstimulated cells, indicating that the compound opposes the loss of receptor from the surface rather than simply adding to the receptor pool **(Fig. 5A)**.

The most dramatic effect was observed with the IFN3 receptor subunit, IL10Rβ **(Fig. 5A)**. MG132 treatment increased surface IL10Rβ in unstimulated cells by ∼1.5-fold within 1 h (P < 0.0001), that is sustained at 6 h. Unlike IFN1, ligand stimulation did not induce substantial degradation of IL10Rβ, consistent with IFN3’s prolonged signaling profile compared to IFN1 ^27^. As a result, increasing IL10Rβ surface expression allows increased signaling throughout the treatment time course from the increased receptor pool. These results indicate that for IFN3, IL10Rβ availability at the cell surface is a rate-limiting step for maximizing signal transduction. The increase in surface receptor was not the result of increased transcription, as absolute transcript abundance for IFNAR1, IFNAR2, IFNλR1, and IL10Rβ transcript concentrations were indistinguishable between MG132 and control cells, regardless of IFN stimulation **(Fig. 5B)**.

## DISCUSSION

As mediators of nearly all immune and inflammatory processes, cytokines are essential for maintaining good health and underlie both acute and chronic diseases. Type I and Type III interferons are the primary cytokines that confer and regulate antiviral immunity ^28^. Both IFNs stimulate activation of JAK-STAT1/STAT2 signaling to induce interferon gene expression and biological responses, but the epithelial-restricted receptor expression of IFN3s has an added benefit in that they do not elicit a strong inflammatory response ^29^, contrasting with IFN1s ^30^, which makes them attractive candidates for therapeutic augmentation of antiviral immunity at mucosal surfaces ^11^. However, IFN3 responses are intrinsically constrained at the receptor/kinase level, limiting maximal STAT1/2 activation and subsequent antiviral efficacy ^5,19^. We reasoned that small molecules targeting these regulatory nodes could provide a pharmacological strategy to enhance IFN3 activity, which allowed us to home in on molecules that act proximal to receptor activation.

Our screening assay reflects key conceptual choices that distinguish it from prior approaches ^31,32^. By measuring pSTAT1 as a receptor-proximal signaling event before downstream transcriptional amplification, we skewed toward identifying compounds that affect upstream regulatory mechanisms. The simultaneous measurement of IFN1 and IFN3 responses on the same plate provides a built-in selectivity filter, enabling direct classification of compounds by cytokine specificity. The Z’ factor ≥ 0.65 confirms assay robustness, and the recovery of known JAK inhibitors as pSTAT1 suppressors validates the platform’s mechanistic sensitivity. Together, these features make the assay a powerful and versatile tool for identifying IFN signaling modulators and allowed the unprecedented identification of previously unrecognized small molecules that can potentiate IFN3 signaling and antiviral function.

The identification of proteasome inhibitors as IFN3 pSTAT activators is a significant finding for several reasons. First, this class of compounds has not been previously associated with IFN3 signaling, defining a new intrinsic regulatory node in the IFN3 system. Despite IFN1 and IFN3 sharing the JAK1/TYK2 kinase machinery and STAT1/STAT2 substrates, proteasome inhibition has a unique ability to enhance IFN3 responses compared with minimal effects on IFN1 signaling across two cell lines and an extended time course. This selectivity implies that the compounds are not acting as broad JAK activators or phosphatase inhibitors but rather are interfering with a previously unappreciated regulatory mechanism that differentially constrains IFN3 versus IFN1 receptor complexes. Second, the functional translation of enhanced STAT phosphorylation into augmented antiviral activity in a VSV-GFP infection assay establishes that the biochemical effect is biologically meaningful. Third, the IFN3 enhancement is rapid (significantly different within 30 minutes) and sustained over time, features that are amenable to use in “tuning” IFN responses for specific outcomes or pathogens.

Proteasome inhibitors are established compounds repurposed here in a new biological context ^33,34^, consistent with the emerging paradigm of phenotypic screening revealing unexpected activities for characterized molecules. Activity at the receptor-proximal step of the pathway offers a coherent and rational explanation for the signaling and antiviral phenotypes. By increasing the number of IL10Rβ chains available at the surface, MG132 raises the ceiling on functional IFNλR1-IL10Rβ heterodimer assembly. IFN3 signaling is limited in part by the low surface abundance of its receptor, so an increase in the limiting subunit would amplify JAK1/TYK2 recruitment and STAT1/STAT2 phosphorylation, in agreement with our findings. Smaller increases in IFNAR2 occur without a commensurate increase in IFN1 signaling ^35^, consistent with the IFN1 response operating near saturation in these cells, where receptor abundance is not rate-limiting ^19^. Another key observation is that the IFN3 receptor chains are not susceptible to ligand-induced degradation, providing a plausible explanation for the well-documented IFN3 prolonged signaling profile compared to the transient kinetics of IFN1 signaling.

Modulation of IFN3 receptor subunit stability thus defines a new regulatory node that can be engaged pharmacologically to selectively potentiate IFN3 signaling. The therapeutic implications of this work are significant, as respiratory and enteric viruses, including notable biomedical concerns like SARS-CoV-2, influenza, norovirus, and respiratory syncytial virus, are primary targets for IFN3-based interventions given their replication at epithelial barriers ^11^. Clinical trials of pegylated IFN3 have demonstrated antiviral activity with a substantially improved tolerability profile compared to IFN1 ^36–39^.A small molecule that potentiates endogenous IFN3 signaling or enhances the activity of administered IFN3 could offer a complementary or standalone antiviral strategy with favorable pharmacological properties. Available proteasome inhibitors, or next-generation analogs developed through medicinal chemistry, represent candidate starting points for such therapeutic development.

Despite the novelty of these findings, important questions remain, including the extent to which proteasome inhibitors enhance IFN3 activity in physiologically relevant organismal contexts and infection models. The generality of enhancement across IFN3 family members and against diverse RNA and DNA viruses requires additional assessment, but all IFN-sensitive infections will likely be similarly affected. It is tantalizing to speculate that other cytokine receptor subunits may also be sensitized by proteasome inhibitors. The HTS platform described here could be readily adapted for other cytokine JAK-STAT pathways, and expansion of our screen to larger compound collections will continue to yield novel modulators that will help address these questions and further define the mechanisms of IFN3 signaling.

In conclusion, we have established a validated high-throughput platform for discovery of IFN3 signaling modulators and identified a family of small molecules that specifically enhance IFN3-mediated STAT activation and antiviral immunity. These compounds represent new chemical tools for probing IFN3 biology and a foundation for therapeutic development aimed at enhancing mucosal antiviral defenses

## MATERIALS AND METHODS

### Cell lines and culture

A549 (human lung adenocarcinoma) and HEK293T (human embryonic kidney) cells were obtained from ATCC and maintained in Dulbecco’s modified Eagle’s medium (DMEM) supplemented with 10% fetal bovine serum (FBS) and 1% penicillin-streptomycin at 37°C with 5% CO_2_. All cell lines were authenticated by STR profiling and confirmed mycoplasma-negative before use.

### Cytokines and compounds

Recombinant human IFNα (referred to as IFN1) was obtained from Hoffmann-La Roche Inc. (product identifier C122643-02, Nutley, NJ, USA); recombinant human IFNω, IFN3, and IFN3 H11 (IFNλ3-H11 variant) were recombinantly expressed and purified as previously described (Mendoza et. al.). Compounds CPD1–6 were obtained from MedChem Express (for Bortezomib, Cat# HY-10227, Epoxymyocin, Cat# HY-13821, and Ixazomib, Cat# HY-10453), Sigma-Aldrich (for MG-132, Cat# 474790), and Neta Scientific (for Delanzomib, Cat# CAYM-23557-1, Oprozomib, Cat# CAYM-16269-1) and dissolved in dimethylsulfoxide (DMSO) to prepare stock solutions at 10 mM. Working concentrations were prepared by serial dilution in culture medium. Final DMSO concentration did not exceed 0.1% in any experiment. Vehicle controls used matched DMSO concentrations.

### In-cell western (ICW) assay for pSTAT1 and pSTAT2

The ICW assay was developed and optimized in collaboration with the Northwestern University High Throughput Analysis Laboratory (HTAL). Briefly, cells were seeded in 96- or 384-well plates at optimized densities (10,000 cells/well for 96-well; 5000 cells/well for 384-well) and allowed to adhere overnight. Compounds were diluted in assay medium and added to cells for 60 minutes before IFN stimulation. Cells were then stimulated with IFN1 or IFN3 at [diluted 1:1000] units/mL for 60 minutes at 37°C. Following stimulation, medium was removed, and cells were fixed with 1.5% paraformaldehyde for 15 minutes at room temperature, permeabilized with ice-cold methanol for 5 minutes at -80°C, and blocked with 10% goat serum, 1% BSA, and 0.1% Tween-20 in PBS for 1 hour. Cells were incubated with primary antibodies against pSTAT1 Y701 (Cell Signaling, Cat# 7649S; 1:1000) and pSTAT2 Y689 (Cell Signaling, Cat# 88410S; 1:1000) overnight at 4°C, washed, and incubated with HRP secondary antibodies. Luminescence was developed using the Pierce SuperSignal ELISA femto reagent and quantified using a BioTek Synergy Neo2 plate reader. Assay quality was assessed using the Z’ factor calculated from on-plate positive (IFN only, no compound) and negative (no IFN, no compound) controls.

### High-throughput compound screening

Six compound libraries representing ∼8,640 compounds were screened at the NU HTAL using the optimized ICW pSTAT1 assay in 384-well format. Compounds were tested at a single concentration (10 μM) with 0.1% DMSO vehicle control. Each compound was tested in the presence of IFN1 and IFN3 simultaneously, with unstimulated vehicle wells and IFN-only vehicle wells as on-plate controls. Raw fluorescence values were normalized to on-plate controls and expressed as a percent of IFN-only response. Hit compounds were defined as those increasing pSTAT1 by >20% over IFN-only controls in at least one IFN condition without >30% increase in cytotoxicity (assessed by Cell-Titer Glo assay, Promega). Hits were counter-screened by dose-response (4-point, 0.1-10uM, in triplicate) in both IFN1 and IFN3 plus no IFN conditions.

### Immunoblot analysis

Cells were stimulated as described and harvested by first washing with cold phosphate-buffered saline and then lysed with whole-cell extract buffer (WCEB) consisting of 50 mM Tris, 280 mM NaCl, 0.5% NP-40, 0.2 mM EDTA, 2 mM EGTA, 10% glycerol, 1 mM dithiothreitol (DTT), 2.5 mM sodium vanadate, and protease inhibitors. Lysates were resolved by SDS-PAGE, transferred to nitrocellulose membranes, and probed with commercial primary antibodies recognizing phospho STAT1 Y701 (Cell Signaling, Cat# 7649S at 1:4,000), phoshpho STAT2 Y690 (Cell Signaling, Cat# 88410S at 1:2,000), total STAT1-C (Santa Cruz Biotechnology, Cat# sc-345 at 1:2,000) and total STAT2-C (Santa Cruz Biotechnology, Cat# sc-476 at 1:1,000). Proteins were visualized by enhanced chemiluminescence (PerkinElmer, catalog no. NEL105001EA) and imaged using a UVP BioSpectrum Imaging System. Digital files were exported to Adobe Photoshop and Illustrator for cropping and figure assembly. Intact digital files are only subjected to minor contrast and quality modifications applied to the entire image and shown intact without lane splicing.

### Antiviral activity assay (VSV-GFP infection)

A549 cells were plated at 2x10^5^ in chamber slides (Thermo Fisher cat# 177445). The following day, cells were pre-treated with compounds at 10μM together with IFNs at 1nM for 4 hours and then infected with VSV-GFP (Indiana strain) at an MOI of 0.1 for 16 hours. Fluorescent images were obtained 24h post infection using a Nikon Eclipse Ti fluorescence microscope equipped with a Photometrics CoolSnap CF2 camera at 10x magnification. Images were adjusted for brightness and contrast using Adobe Photoshop.

### Cell Surface Receptor Expression

A549 cells (from Ola) were maintained in high-glucose DMEM supplemented with GlutaMAX™ (Gibco/Thermo Fisher, 10569-044), 10% fetal bovine serum (FBS; Corning, 35-010-CV), and 1% penicillin–streptomycin (Cytiva, SV30010), at 37°C with 5% CO_2_. Cells were seeded at 30,000 cells per well in 100 µL culture medium in tissue culture–treated, flat-bottom 96-well plates (Celltreat, 229196) and cultured overnight.

For MG-132 and interferon (IFN) treatment, culture medium was removed and replaced with 50 µL medium containing 10 µM MG132 (Sigma-Aldrich, 474790); cells were incubated for 1 h at 37°C with 5% CO_2_. Following incubation, 10 µL of IFN was added to designated wells to achieve final concentrations of 1 nM IFN-ω1 (type I IFN) or 100 nM IFNλ3 (type III IFN), both lab-generated. Cells were incubated for an additional 0, 1, or 6 h at 37°C with 5% CO_2_.

Following treatment, culture medium was removed, and cells were incubated with 50 µL TrypLE Express (Gibco/Thermo Fisher, 12605010) per well for 5–10 min at 37°C to detach cells. Cells were gently resuspended in 100 µL culture medium, centrifuged at 1500rpm/520xg (Beckman Coulter, Avanti-15R with JS -4.750 rotor) for 4 min, and the supernatant was removed. Cells were washed twice with 150 µL PBS containing 0.5% bovine serum albumin (PBSA, Thermo Fisher, AAJ6509722), with centrifugation as above between washes.

For surface receptor staining, cells were resuspended in 20 µL antibody staining solution and incubated for 1 h on ice. Antibodies were used at the manufacturers’ recommended dilutions. Surface expression of IFNAR1, IFNAR2, and IL10Rβ was assessed directly using PE-conjugated primary antibodies, whereas IFNLR1 was detected using an unconjugated primary antibody followed by a PE-conjugated secondary antibody. PE-conjugated anti-transferrin receptor staining was performed in parallel as a control. For IFNλR1 staining, cells were washed twice following primary antibody incubation and then incubated with PE-conjugated secondary antibody for 30 min on ice.

### Cell Culture and Preparation

A549 cells were cultured in DMEM supplemented with 10% fetal bovine serum (FBS) and 1% penicillin-streptomycin (P/S) at 37°C and 5% CO_2_. Cells were seeded at 1.5x10^5^ cells/well in 6-well plates one day prior to treatment. Cells were assigned to six experimental conditions: untreated control, IFNω, IFNω + MG132, IFNλ3, IFNλ3 + MG132, and MG132 only. Cells receiving MG132 were pretreated with 10 μM MG132 for 1 h, followed by the addition of the respective interferon. Cells were then incubated for 24 h at 37°C before downstream analysis.

### RNA Extraction and cDNA Synthesis

Total RNA was extracted using the Monarch Total RNA Miniprep Kit (T2010) according to the manufacturer’s protocol. RNA concentration was measured using the Qubit™ RNA Broad Range (BR) Assay Kit (Q10210). 1 µg of total RNA was then reverse-transcribed into cDNA using the Applied Biosystems High-Capacity cDNA Reverse Transcription Kit (4368814) according to the manufacturer’s protocol, and diluted with 60 µL of water.

### Digital Polymerase Chain Reaction (dPCR)

Digital polymerase chain reaction (dPCR) was performed to quantify absolute transcript concentrations using the Absolute Q Universal DNA Digital PCR Master Mix (5X) (A72710) and QuantStudio Absolute Q MAP16 Plate Kit (A52865) according to the manufacturer’s protocol. Gene expression was assessed using FAM-MGB–labeled TaqMan Gene Expression Assays specific for IFNλR1 (Hs00417120_m1), IL10Rβ (Hs00175123_m1), IFNAR1 (Hs01066116_m1), and IFNAR2 (Hs01022059_m1). Following amplification, absolute target concentrations were determined and reported as copies per microliter (copies/µL).

### Statistical analysis

Statistical analyses were performed using GraphPad Prism version 11. Comparisons between two groups were made using an unpaired t-test / Mann-Whitney U test. Multiple group comparisons were made using one-way ANOVA with Tukey’s post-hoc test. All experiments were performed in at least three independent biological replicates unless otherwise noted. Data are represented as mean ± SD or mean ± SEM as indicated in figure legends. P values < 0.05 were considered statistically significant.

## ACKNOWLEDGMENTS

We thank the Northwestern University High Throughput Analysis Laboratory (HTAL) for expert assistance with assay development and screening. This work was supported by the Chicago Biomedical Consortium to J.L.M. and C.H.. Funding to support J.L.M. is provided by the Pritzker School of Molecular Engineering at the University of Chicago, the National Institutes of Health (NIH) grant 1R35GM147179-01, and the Howard Hughes Medical Institute.

## AUTHOR CONTRIBUTIONS

All authors contributed to writing and reviewing the manuscript. J.L.M. conceived of the idea for a screen. J.L.M. and C.H. designed the approach and oversaw experiments. S.F. oversaw the HT screen. J-P.P. performed signaling and AV assays. S.S. and L.T.H. performed dPCR experiments. C.O’L. performed receptor surface experiments. A.K. performed preliminary signaling studies.

---

**Table 1.** Antibodies used for flow cytometry.

| Antibody | Manufacturer | Catalog# | Usage |
| --- | --- | --- | --- |
| PE-IFNAR1 | Invitrogen/ThermoFisher | MA5-23630 | Add 10 μL per well |
| PE-IFNAR2 | SinoBiologicals | 10359-R122-P | 1:50 dilution in 20 μL |
| Unconjugated IFNLR1 | OriGene/ThermoFisher | TA812609S | 1:100 dilution in 20 μL |
| Secondary PE | Invitrogen/ThermoFisher | P-852 | 1:100 dilution in 20 μL |
| IL10RB | Invitrogen/ThermoFisher | MA5-50604 | 1:20 dilution in 20 μL |
| Transferrin R | Invitrogen/ThermoFisher | MHCD7104 | Add 10 μL per well |

